# METTL3 alters capping enzyme expression and its activity on ribosomal proteins

**DOI:** 10.1101/2023.11.22.568301

**Authors:** Daniel del Valle-Morales, Giulia Romano, Patricia Le, Michela Saviana, Rachel Brown, Lavender Micalo, Howard Li, Alessandro La Ferlita, Giovanni Nigita, Patrick Nana-Sinkam, Mario Acunzo

## Abstract

The 5’ cap, catalyzed by RNA guanylyltransferase and 5’-phosphatase (RNGTT), is a vital mRNA modification for the functionality of mRNAs. mRNA capping occurs in the nucleus for the maturation of the functional mRNA and in the cytoplasm for fine-tuning gene expression. Given the fundamental importance of RNGTT in mRNA maturation and expression there is a need to further investigate the regulation of RNGTT. N6-methyladenosine (m^6^A) is one of the most abundant RNA modifications involved in the regulation of protein translation, mRNA stability, splicing, and export. We sought to investigate whether m^6^A could regulate the expression and activity of RNGTT. A motif for the m^6^A writer methyltransferase 3 (METTL3) in the 3’UTR of RNGTT mRNA was identified. Knockdown of METTL3 resulted in destabilizing RNGTT mRNA, and reduced protein expression. Sequencing of capped mRNAs identified an underrepresentation of ribosomal protein mRNA overlapping with 5’ terminal oligopyrimidine (TOP) mRNAs and genes are dysregulated when cytoplasmic capping is inhibited. Pathway analysis identified disruptions in the mTOR and p70S6K pathways. A reduction in RPS6 mRNA capping, protein expression, and phosphorylation was detected with METTL3 knockdown.

## Introduction

The 5’ cap, which consists of an N^7^-methylated guanosine molecule, is covalently added to the first transcribed nucleotide by the mRNA capping enzyme RNGTT [1]. The 5’ cap coordinates the downstream functions of mRNAs, and is required for protein translation, mRNA stability, splicing, and export [2]. The 5’ cap is canonically added in the nucleus to all RNAs transcribed by RNA Polymerase II [3]. In the nucleus, RNGTT along with RNA guanine-7 methyltransferase (RNMT) [4] and its coactivator RNA guanine-7 methyltransferase activating subunit (RAMAC) [5] interacts with the C-terminal domain of RNA Polymerase II to add the 5’ cap during the initial stages of transcription [6]. mRNA capping is also detected in the cytoplasm [7] where RNGTT, RNMT-RAMAC, and an unknown 5’monophosphatase is bound to the scaffold protein NCK1 [8, 9]. In the cytoplasm, the cap of uncapped mRNAs [10] and long noncoding RNAs [11] are restored by the cytoplasmic capping complex, serving as a regulatory mechanism to fine-tune gene expression [10].

In addition to its basic function in mRNA maturity, RNGTT has been linked to biological and pathological processes such as development [12] and certain cancers [13]. RNGTT was shown to positively modulate the Hedgehog signaling pathway by inhibiting PKA kinase activity in *Drosophila* development. The mammalian homolog of RNGTT was shown to have the same function in *Drosophila* and cultured mammalian cells [12]. The oncogene c-MYC recruits RNGTT to RNA Polymerase II during transcription; the cell proliferation of c-MYC overexpressing cells is reduced with the downregulation of RNGTT while cells with normal c-MYC levels were unaffected [14]. Furthermore, RNGTT was upregulated in high-eIF4E acute leukemia patients and increased mRNA capping of MYC, CDK2, and MALAT1 was detected [15]. Given the observed importance of RNGTT, there is a need to better understand regulatory pathways of mRNA capping in order to investigate other biological functions of RNGTT.

A possible regulatory pathway for capping involves epitranscriptomic modifications of RNGTT mRNA such as m^6^A. m^6^A, one of the most abundant modifications in RNAs [16], is a dynamic marker in RNAs that is added by m^6^A writers, removed by m^6^A erasers, and recognized by m^6^A readers to undergo its functional role [17]. m^6^A modification in mRNAs has been associated with enhanced protein translation, mRNA stability, splicing, export [17], and linked to a wide range of biological processes [17].

In this study, we identified and validated an m^6^A site in the 3’UTR of RNGTT mRNA. Using non-small cell lung cancer (NSCLC) cell lines as our model, we demonstrated a regulatory pathway where METTL3 directly methylates this region and affects the RNA stability and protein expression of RNGTT. We identified the disruption of mRNA capping on ribosomal protein mRNAs, and alterations of the mTOR and p70S6K signaling pathways.

## Results and Discussion

### m^6^A modification of RNGTT 3’UTR by METTL3 regulates its protein expression and RNA stability

Using data mining on the RNA EPItranscriptome Collection (REPIC) [18], we identified a region at the 3’UTR of RNGTT enriched in m^6^A pulldowns. The 3’UTR of RNGTT was then scanned for the consensus m^6^A motif DRACH [16] using the FIMO motif scan tool [19, 20]. Both datasets are represented as a genome track (Fig. 1A left). The m^6^A motifs that were identified with FIMO are represented in Fig. 1A right. Taken together, these sites predict m^6^A modification sites at the 3’UTR of RNGTT. To study the potential role of this m^6^A site of RNGTT, we generated A549 cells that stably express either a scramble shRNA (sh-) or shMETTL3. We performed western blot analysis probing for METTL3 and RNGTT and observed the reduction of METTL3 in shMETTL3 cells (Fig. 1B). RNGTT protein expression was reduced in shMETTL3 cells when compared to sh-cells (Fig. 1B). Additionally, we verified that shMETTL3 cells have reduced global m^6^A levels compared to sh-as expected (Fig. 1C). To confirm that METTL3 directly methylates RNGTT, m^6^A RNA immunoprecipitation followed by RT-qPCR was performed using primers for the predicted m^6^A region in RNGTT mRNA (Fig. 1D left). The probed region of RNGTT was reduced in shMETTL3 compared to sh-after m^6^A immunoprecipitation (Fig. 1D right), confirming that METTL3 directly methylates RNGTT mRNA in the predicted site.

**Figure 1.**
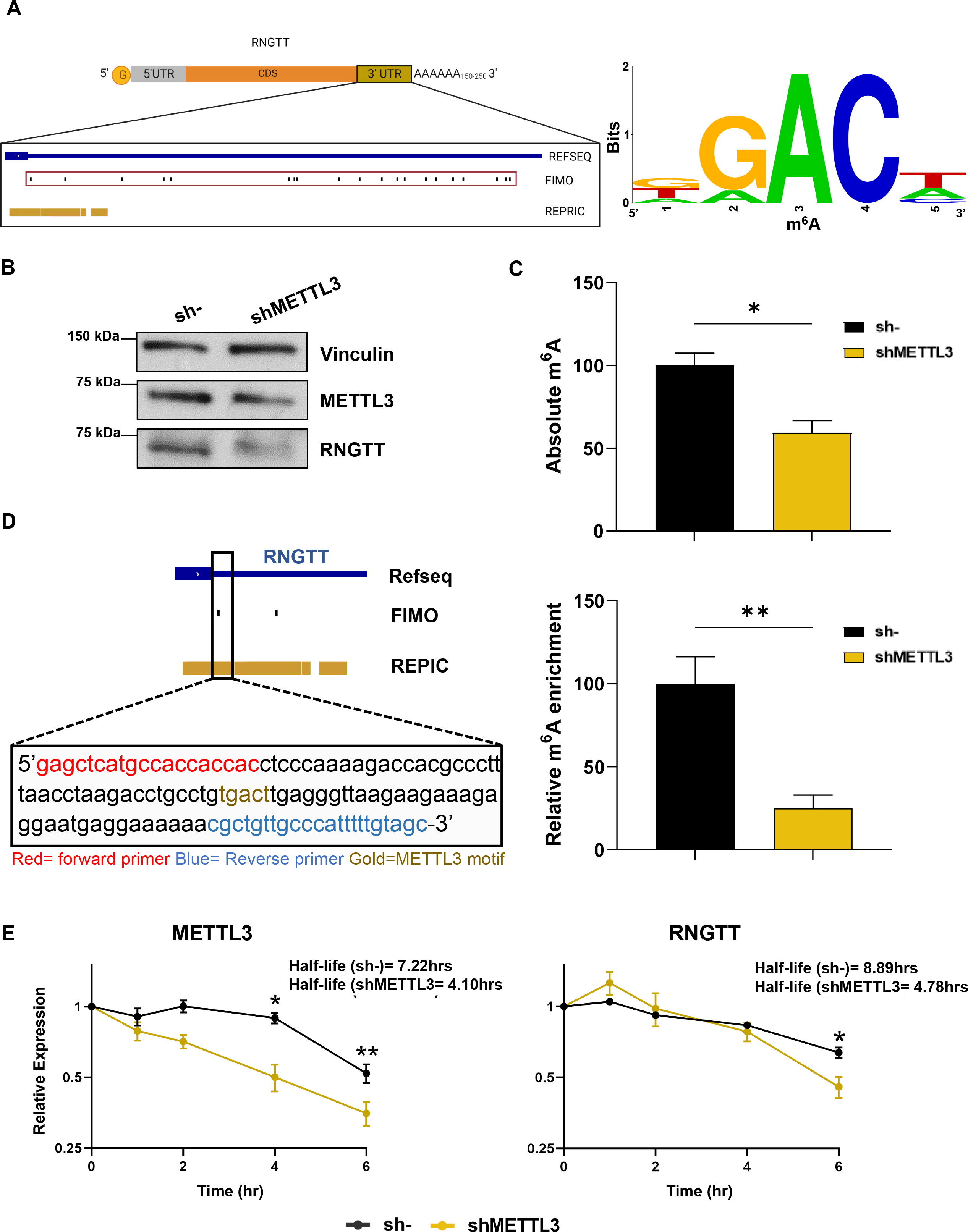
METTL3 regulates the expression and stability of RNGTT. A) Genomic view of the 3’UTR of RNGTT. m^6^A pulldown peaks (REPRIC) and METTL3 motif prediction (FIMO) are displayed as tracks in IgvR (left). The METTL3 motif prediction tracks (red box) are represented as a sequence logo (right). B) Representative western blot analysis of protein expression of METTL3, RNGTT, and loading control Vinculin done in triplicate. C) Global m^6^A levels of A549 sh-/shMETTL3. m^6^A levels are represented as absolute m^6^A levels with sh-set as 100%. Error bars represent S.E.M. in technical triplicate n=3. * represents two-tailed student t-test of sh-vs shMETTL3 p. value <0.05. D) m^6^A RNA immunoprecipitation on predicted site of m^6^A modification of RNGTT mRNA analyzed by RT-qPCR. Immunoprecipitation is represented as % input relative to recovery of m^6^A positive control. sh-was set to 100%. Error bars represent S.E.M. in six biological replicates N=6. ** represents two-tailed student t-test p. value <0.01. E) RNA stability assay in A549 sh-/shMETTL3 analyzed by RT-qPCR. RNA expression is represented as RNA levels normalized to GAPDH at the indicated timepoint relative to the RNA levels at timepoint 0 hours in either sh- or shMETTL3. Error bars represent S.E.M. of biological triplicates N=3. * represents two-tailed student t-test of sh-vs shMETTL3 p. value <0.05 at each indicated timepoint, ** represents p. value <0.01.

To determine the mechanism by which METTL3 reduces RNGTT protein expression, we examined the RNA stability of RNGTT in sh- and shMETTL3 cells. sh- and shMETTL3 cells were treated in triplicate in a time course with the RNA Polymerase II inhibitor 5,6-dichlorobenzimidazole (DRB) [21]. The DRB time course showed the expected reduction of the RNA stability of METTL3 mRNA (sh-RNA half-life=7.22hrs, shMETTL3 RNA half-life=4.10hrs) (Fig. 1E left). The RNA stability of RNGTT is significantly reduced in shMETTL3 cells at the 6hr timepoint (sh-RNA half-life= 8.89hrs, shMETTL3 RNA half-life=4.78hrs), suggesting that the m^6^A site on RNGTT increases the RNA stability of RNGTT (Fig. 1E right). Our findings do not exclude the possibility that other targets of METTL3 could affect the protein expression and mRNA stability of RNGTT, in addition to the reported effects by the m^6^A siteMETTL3 itself has been shown to increase the protein translation of mRNAs independently of its m^6^A methylation activity (30).

### Knockdown of METTL3 reduced the RNA capping of ribosomal proteins

The observed decreased of RNGTT with METTL3 knockdown raises the question of whether global mRNA capping would also be altered. Uncapped mRNA are translationally inactive and are subject to 5’ RNA degradation [2] or are stored in an uncapped state [10]. To test the global changes in capped mRNAs, we performed capped cDNA enrichment using TeloPrime 5’ cDNA kit (Lexogen) followed by next-generation sequencing in A549 sh- and shMETTL3 A549 cells (Fig. 2A). TeloPrime enriches for capped mRNAs through double strand ligation of an adaptor with a cytosine overhang to the 5’ guanosine cap. Through PCR amplification of the adapter, capped mRNAs are amplified while uncapped mRNAs are excluded. The resulting cDNA represents the capped population of mRNAs which allows the precise analysis of differentially capped mRNAs. Interestingly, differential expression analysis of sh-vs shMETTL3 A549 cells identified a subset of 416 dysregulated genes (≥ 1.5-fold change and p.val. <0.001), despite the observed downregulation of RNGTT protein expression. The differentially capped mRNAs were distributed between 126 downregulated and 298 upregulated genes (Fig. 2B and Table S1).

**Figure 2.**
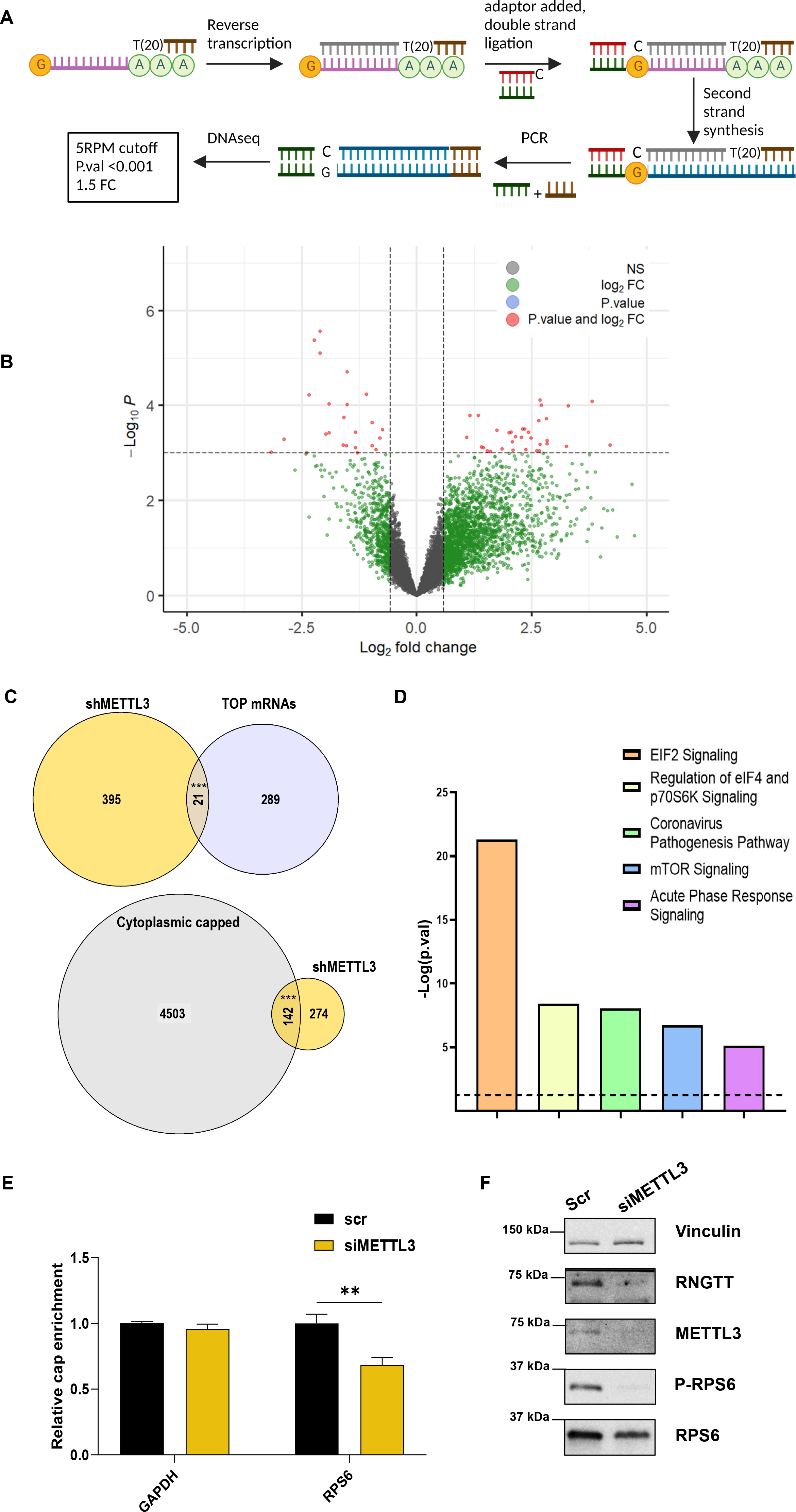
Downregulation of METTL3 disrupts mRNA capping in select genes. A) Schema of TeloPrime capped mRNA cDNA synthesis and DNAseq analysis. B) Volcano plot on differentially expressed genes sh-/shMETTL3 A549. Grey points represent no significance (p. value >0.001 and |Log2FC| < 0.58), green points represent |Log2FC| > 0.58, blue points represent p-value < 0.001), and red points represent significant genes (|Log2FC| > 0.58 and a p-value < 0.001). C) Gene overlap between top mRNAs or cytoplasmic capped mRNAs and differentially expressed genes. Venn diagrams are scaled with list size. *** represents overlapping Fisher exact test p. value <0.001. D) Histogram of the top 5 pathways identified by IPA for the downregulated capping genes. Dotted line represents the threshold for p. value <0.05. E) CAP-IP on selected genes in PC9 cells analyzed by RT-qPCR. Data were first normalized to input and then normalized to housekeeping control ACTIN and represented as fold change with scr enrichment set as 1. Error bars represent S.E.M. in N=7.. Ns represents non-significant two-tailed student t-test of scr vs siMETTL3. * represents two-tailed student t-test of scr vs siMETTL3 p. value <0.05, ** represents p. value <0.01. F) Representative western blot analysis of protein expression of METTL3, RNGTT, P-RPS6, RPS6 and Ponceau staining as loading control done in triplicate.

To expand the observed relation between METTL3 and RNGTT, we transiently repressed METTL3 in an additional NSCLC cell line, PC9 (Fig. S1). We observed a decrease in RNGTT protein expression (Fig. S1A), m^6^A pulldown (Fig. S1B), and mRNA stability in PC9 cells (scr-RNA half-life=7.22hrs, siMETTL3 RNA half-life=4.10hrs) (Fig. S1C) as was observed in the A549 cell line. Furthermore, we performed CAP-IP [15] to assay the difference in the capping status of these genes when METTL3 is downregulated as an independent validation of the sequencing data. We focused on a selected set of downregulated ribosomal proteins, such as RPL7, RPL29, and RPL13A (all with a fold change <-1.5). We observed a significant decrease of CAP-IP fold enrichment for RPL7, and RPL29 with no change in housekeeping gene GAPDH in siMETTL3 transfected PC9 cells (Fig. S1D). There was a decreasing trend of RPL13A, which was not statistically significant. Thus, the decrease in CAP-IP enrichment enforces the changes in differentially capped mRNAs observed in A549 cells.

To identify the class of genes that were differentially capped in our sequencing data (Figure 2B), we performed a protein class analysis through the PANTHER tool [22, 23] which identified translational and ribosomal proteins as underrepresented and scaffold/adaptor proteins as overrepresented in our differentially capped mRNAs (Table S2). A previous study identified a downregulation of TOP mRNAs, which constitute translational and ribosomal proteins [24], when cytoplasmic capping is inhibited [25].

We explored whether our differentially capped mRNAs shared similarities with known cytoplasmic capping targets. TOP mRNAs are translationally regulated by La-related protein 1 (LARP1) which binds to both the TOP motif and the 5’cap [26, 27]. We examined if there was significant overlap between TOP mRNAs identified using LARP1 RNA Immunoprecipitation performed in Gentilella et. al. [28] and our differentially capped mRNAs, and found a significant overlap of 21 genes between the two lists (Fig. 2C top and Table S3). When examining the cytoplasmic capped transcripts [25], we observed a significant overlap of 142 genes (Fig. 2C bottom and Table S3). This data suggests that METTL3 can affect cytoplasmic capped mRNAs and METTL3-mediated methylation could be affecting the recruitment of uncapped mRNAs to the cytoplasmic capping complex. RNGTT has been observed to be distributed into a nuclear and cytoplasmic population [7]. The downregulation of RNGTT protein may also induce a shift in the sub-population of RNGTT where the canonical nuclear population is retained or increased while the cytoplasmic population of RNGTT is reduced, which may explain why only a small subset of capped mRNAs was affected with METTL3 knockdown.

To determine the possible functional role of the differentially capped mRNAs, we performed a gene set analysis through Ingenuity Pathway Analysis (IPA) for all genes and downregulated capped genes (Table S4). We focused on the pathways identified with the downregulated capped genes as these are more directly linked to the reduction of RNGTT. The top five significant canonical pathways of the downregulated capped genes are represented in Figure 2D. As alluded to from the gene overlap of Figure 2C, three of the five pathways identified are involved in protein translation, in particular, p70S6K signaling and the mTOR signaling pathway. Interestingly, p70S6K is downstream the mTOR pathway [29] and directly phosphorylates RPS6, a marker of the activation of both pathways [30, 31]. When we examined the capping status of RPS6 in the PC9 cap pulldown, there was a significant decrease of RPS6 capping (Figure 2E) suggesting that RPS6 signaling may be impaired with METTL3 knockdown. To test for the reduction of these two pathways, we performed western blot on the phosphorylation of RPS6 with the knockdown of METTL3 A549 cells. We observed a reduction of P-RPS6 signal with siMETTL3 and a reduction of total RPS6 in line with our differential capping analysis and the CAP-IP (Figure 2F). METTL3 has been shown to be a positive regulator for both the mTOR and p70S6K [32, 33]. The reduced capping of genes in these pathways suggests that METTL3 might contribute to the activation of mTOR and p70S6K through RNGTT.

Taken together, the knockdown of siMETTL3 reduces the RNA capping of ribosomal proteins and impacts the phosphorylation of a downstream target of the mTOR and p70S6K signaling pathway.

## Conclusions

We identified METTL3 as a regulator of RNGTT mRNA stability and protein expression in NSCLC cell lines through the m^6^A methylation at the 3’UTR. Interestingly, downregulation of METTL3 altered the RNA capping of a small set of mRNAs which overlapped with TOP mRNAs and cytoplasmic capped mRNAs. We found and validated the decrease in capping of RPL7, and RPL29. Functionally, the downregulated capped mRNAs were associated with mTOR and p70S6K signaling. We observed a reduction in the mRNA capping of RPS6 and P-RPS6 with METTL3 knockdown confirming the alteration of the mTOR and p70S6K signaling pathway. These pathways are critical for several physiological and pathological mechanisms [34], thus, the regulatory relationship between METTL3 and RNGTT should be deeply investigated in other biological context such as disease and cancer.

## Experimental procedures

### Cell culture and cell line development

A549 and PC9 cell lines were seeded and grown in RPMI (Sigma-Aldrich #R8758), supplemented with 10% FBS (Sigma #F4135) and 1% Penicillin/Streptomycin (Quality Biological #120-095-721). Stable A549 cell lines were generated with transfection of sh-(SantaCruz #sc-108060) and shMETTL3 (SantaCruz #sc-92172) using Lipofectamine LTX (ThermoFisher # A12621). Stable cells were selected and grown in RPMI + 0.75μg/mL of puromycin. All cells were synchronized with RPMI + 0.5% FBS prior to cell plating.

### Western blot

Cells for protein extraction were lysed using RIPA lyses buffer (Sigma #R0278) containing protease and phosphatase inhibitor cocktails (cOmplete™, Mini Protease Inhibitor Cocktail, Sigma #11836153001; PhosSTOP™, Roche #4906845001). Protein quantification was determined using the BCA protein assay (Pierce™ BCA Protein Assay Kit, Thermo-Fisher #23225) following the manufacturer’s instructions. Equal amounts of protein lysates and rainbow molecular weight marker (Bio-Rad Laboratories #1610394) were separated by 7.5% SDS/PAGE (Bio-Rad #4561106) and then electro-transferred to 0.2μm nitrocellulose membranes (Bio-Rad # 1620112). The membranes were blocked with a buffer containing 5% nonfat dry milk in Tris-buffered saline with 0.01% Tween20 (TBST) for at least 1hr and incubated overnight with primary antibodies at 4°C (anti-METTL3, ThermoFisher # 15073-1-AP; anti-RNGTT, ThermoFisher # 12430-1-AP; anti-Vinculin, Santa Cruz #sc-25336; anti-GAPDH, Abcam #ab8245; anti-S6 Ribosomal Protein, Cell Signaling #5G10; anti-P-S6 Ribosomal Protein, Cell Signaling, D57.2.2E). After three washes with TBST, the membranes were incubated with peroxidase-conjugated secondary antibodies (Anti-rabbit IgG, HRP-linked Antibody, Cell Signaling #7074; Anti-mouse IgG, HRP-linked Antibody, Cell Signaling #7076) and developed with a luminol-based enhanced chemiluminescence (ECL) horseradish peroxidase (HRP) substrate (SuperSignal West Dura Extended Duration Substrate, Thermo Scientific™ #34076)

### Transfection

200,000 cells were seeded on a 6-well dish for 24hrs before transfection. Transfections were performed using Lipofectamine 3000 (ThermoFisher # L3000015) following manufacture’s protocol. Cells were transfected either with negative control (scr-) (Santa Cruz #sc-37007) or siMETTL3 (ThermoFisher # 4392420). The media was changed after 5hrs and transfected cells were harvested after 48hrs.

Transfection efficiency was determined with reverse transcription of 500ng of RNA using High-Capacity cDNA Reverse Transcription Kit (Applied Biosystems™ #4368814). RT-qPCR was then performed with Taqman® technology using the following probes: Actin (Hs99999903_m1), METTL3 (Hs00219820_m1).

### m^6^A site prediction

m^6^A RNA immunoprecipitation peaks were datamined from the REPRIC web browser (https://repicmod.uchicago.edu/repic/) [18]for A549 cells. MeTPeak for RNGTT were downloaded and converted into Hg38 chromosome coordinates. The 3’UTR of RNGTT was downloaded from Ensembl [35] and scanned for the METTL3 motif DRACH using FIMO [19, 20] with a p-value cutoff of 0.001. Motif locations were converted into Hg38 chromosome coordinates and both the REPRIC and FIMO coordinated were loaded into igvR (v1.18.0) for genome visualization. Sequence logo for the FIMO predicted motifs were generated using WebLogo (https://weblogo.berkeley.edu/) with the default settings [36].

### Global m^6^A quantification

Global m^6^A levels were assayed using EpiQuik m^6^A RNA Methylation Quantification Kit (Colorimetric) (EpigenTek # P-9005-48) following manufacture’s protocol for absolute m^6^A levels with 200ng of total RNA.

### m^6^A RNA immunoprecipitation

7.5ug of total RNA was immunoprecipitated for m^6^A with EpiMark® N6-Methyladenosine Enrichment Kit (NEB) following the manufacture protocol with 10% of the sample RNA taken as input prior to the pulldown. The eluted RNA was suspended with 1mL TRIzol (Invitrogen #15596018) and purified using RNA Clean-Up and Concentration Kit (NORGEN #43200) and eluted with 20μl of H_2_O. 10μl of eluted RNA and input RNA was retro transcribed using High-Capacity cDNA Reverse Transcription Kit (Applied Biosystems™ #4368814). RT-qPCR was performed using SYBR GreenER™ qPCR SuperMix for ABI PRISM™ Instrument (ThermoFisher # 11760100). The primers used are the following: positive m^6^A control (FWD 5 - CGACATTCCTGAGATTCCTGG - 3, REV 5 - TTGAGCAGGTCAGAACACTG – 3’), negative m^6^A control (FWD 5 - GCTTCAACATCACCGTCATTG -3, REV 5 - CACAGAGGCCAGAGATCATTC - 3), RNGTT (FWD 5’-GAGCTCATGCCACCACCAC-3’, REV 5’-GCTACAAAAATGGGCAACAGCG-3’).

### RNA stability assay

RNA stability was assayed through a time-course inhibition of RNA Polymerase II with DRB [21]. 75,000 A549 sh/shMETTL3 or transfected PC9 scr-/siMETTL3 in triplicate were plated into a 24-well plate. 24hrs after plating (48hrs for transfected PC9 cells), cells were treated with 20ug/mL of DRB and incubated at 37C, 5% CO_2_ for the indicated time points (0, 2, 4, 6hrs). After the final timepoint, the media was aspirated from the wells, washed once with PBS, and cells were lysed with 1mL of TRIzol. RNA was purified using RNA Clean-Up and Concentration Kit (NORGEN #43200). 250ng of RNA was retrotranscribed as described above. RT-qPCR was performed with Taqman® technology using the following probes: GAPDH (Hs02758991_g1), Actin (Hs99999903_m1), METTL3 (Hs00219820_m1), and RNGTT (Hs01016932_m1). Half-life measurements were calculated according to Chen et. al. [37].

### TeloPrime

Capped mRNAs were enriched and converted to cDNA with TeloPrime Full-Length cDNA Amplification Kit V2 (Lexogen # 013.24). TeloPrime was perfomed on 2μg of total RNA following the manufacture’s protocol. 18 cycles for endpoint PCR was used for final cDNA amplification for all samples. The resulting cDNA was sent for DNAseq at Genomics Core facility at Virginia Commonwealth University (VCU) and sequenced using NextSeq2000 Sequencer (Illumina).

### Pre-processing

Raw sequencing reads in FASTQ format were quality trimmed, and adapters were removed using Trim Galore (v0.6.6) (https://www.bioinformatics.babraham.ac.uk/projects/trim_galore/). Trimmed reads were then mapped to the human genome (HG38 assembly) using HISAT2 (v2.1.0) [38]. Afterward, the mapped reads in SAM format were converted into BAM format, sorted (for coordinates), and indexed using Samtools (v1.12) [39]. Finally, gene quantification was performed by FeatureCounts (v2.0.0) [40] using Gencode (v39) GTF annotation file.

### Differential Expression Analysis

To perform the differential gene expression (DE) analysis, we first scaled the raw read counts via Read Per Million (RPM) and filtered out low expressed genes, whose geometric mean value was less than five across all samples. Afterward, the raw read counts of the retained genes were log2-transformed leveraging the Voom function and then used for the DE analysis, leveraging the Limma R package (v3.48.3) [41]. Genes with a |Log2FC| > 0.58 (|Linear FC| > 1.5) and a p-value < 0.001 were considered differentially expressed. The volcano plot showing the differentially expressed genes was generated using the EnhancedVolcano R package (v1.10.0).

### Pathway analysis and Gene overlap analysis

Using Ingenuity® Pathway Analysis (IPA®) software (v01-22-01), we performed functional enrichment analysis considering the differentially expressed genes (mentioned above). Settings used for the IPA analysis included experimentally observed data for the human species. For gene overlap, the list of TOP mRNAs (labeled as RP and TOP) were extracted from in Gentilella et. al [28] and the statistically significant cytoplasmic capped mRNAs were extracted from del Valle-Morales et al [25]. Gene overlap was performed using the GeneOveralp R package (v1.34.0) using default parameters. Venn diagrams were generated using the R package VennDiagram (v1.7.3) [42].

### CAP-IP

Capped mRNA was immunoprecipitated following the protocol described in Culjkovic-Kraljacic et. al. [15]. 4μg of total RNA spiked with 1ng/μL of Luciferase Control RNA (Promega #L4561) as a negative uncapped control. 10% of RNA and spike-in was taken as input. Bound RNA was purified as described above. 10μL of bound RNA and 10% input was retro transcribed as described above. RT-qPCR was performed with Taqman® technology using the following probes: Actin (Hs99999903_m1), GAPDH (Hs02758991_g1), RPL7(Hs02596927_g1), RPS6(Hs04195024_g1), RPL13A (Hs04194366_g1), RPL29 (Hs06645107_g1). Pulldown efficiency was determined with RT-qPCR using SYBR GreenER™ qPCR SuperMix for ABI PRISM™ Instrument on Luciferase (primers FWD 5’-CTTATGCATGCGGCCGCATCTAGAGG-3’, REV 5’-CAGTTGCTCTCCAGCGGTTCCATCC-3’).

## Availability of data and materials

NGS data will be deposited into GEO repository prior to publication.

## Supporting information

This article contains supporting information.

## Acknowledgements

The DNA sequencing data included in this study was generated at the Genomics Core facility at Virginia Commonwealth University. Diagrams were created with BioRender.com

## Authors’ contributions

DDVM and MA designed the study. DDVM, GR, PL, LM, MS, RB, AL, and GN coordinated and performed the experiments, and analyzed the corresponding results. DDVM and MA wrote the manuscript. All authors contributed to editing the manuscript.

## Funding and additional information

Project supported by VCU Postdoctoral Independent Research Award (DDVM), American Lung Association (LCDA-922902) (MA), and National Institutes of Health (NIH 1R21CA277525-01) (MA). The content is solely the responsibility of the authors and does not necessarily represent the official views of the National Institutes of Health.

## Figure legend

**Figure S1**

**METTL3 regulates the protein expression and mRNA stability of RNGTT, and the capping of ribosomal proteins in PC9 cells** A) Representative western blot analysis of protein expression of RNGTT and METTL3 with loading control GAPDH done in triplicate. B) m^6^A RNA immunoprecipitation on predicted site of m^6^A modification of RNGTT mRNA analyzed by RT-qPCR. Immunoprecipitation is represented as % input relative to recovery of m^6^A positive control. Scr was set to 1. Error bars represent S.E.M. in six biological replicates N=6. * represents student t-test p. value <0.05. C) RNA stability assay is analyzed by RT-qPCR. RNA expression is represented as RNA levels at the indicated timepoint relative to the RNA levels at timepoint 0 hours in either scr or siMETTL3. Error bars represent S.E.M. of biological triplicates N=3. ** represents student t-test of scr vs siMETTL3 p. value <0.01 at each indicated timepoint, *** represents p. value < 0.001. D) CAP-IP on selected genes in PC9 cells analyzed by RT-qPCR. Data were first normalized to input and then normalized to housekeeping control ACTIN and represented as fold change with scr enrichment set as 1. Error bars represent S.E.M. in N=7. Ns represents non-significant two-tailed student t-test of scr vs siMETTL3. * represents two-tailed student t-test of scr vs siMETTL3 p. value <0.05, ** represents p. value <0.01.

**Table S1**

TeloPrime differentially expressed genes sh-vs shMETTL3 A549.

**Table S2**

PANTHER significant enrichment of protein class in differentially expressed genes

**Table S3**

Gene overlap between TOP mRNAs and cytoplasmic capped mRNAs with differentially expressed genes

**Table S4**

IPA canonical pathway analysis of all dysregulated capped mRNAs and downregulated capped mRNAs

