## Supplementary figures and images for "METTL3 alters capping enzyme expression and its activity on ribosomal proteins"

### Supplemental Figure 1

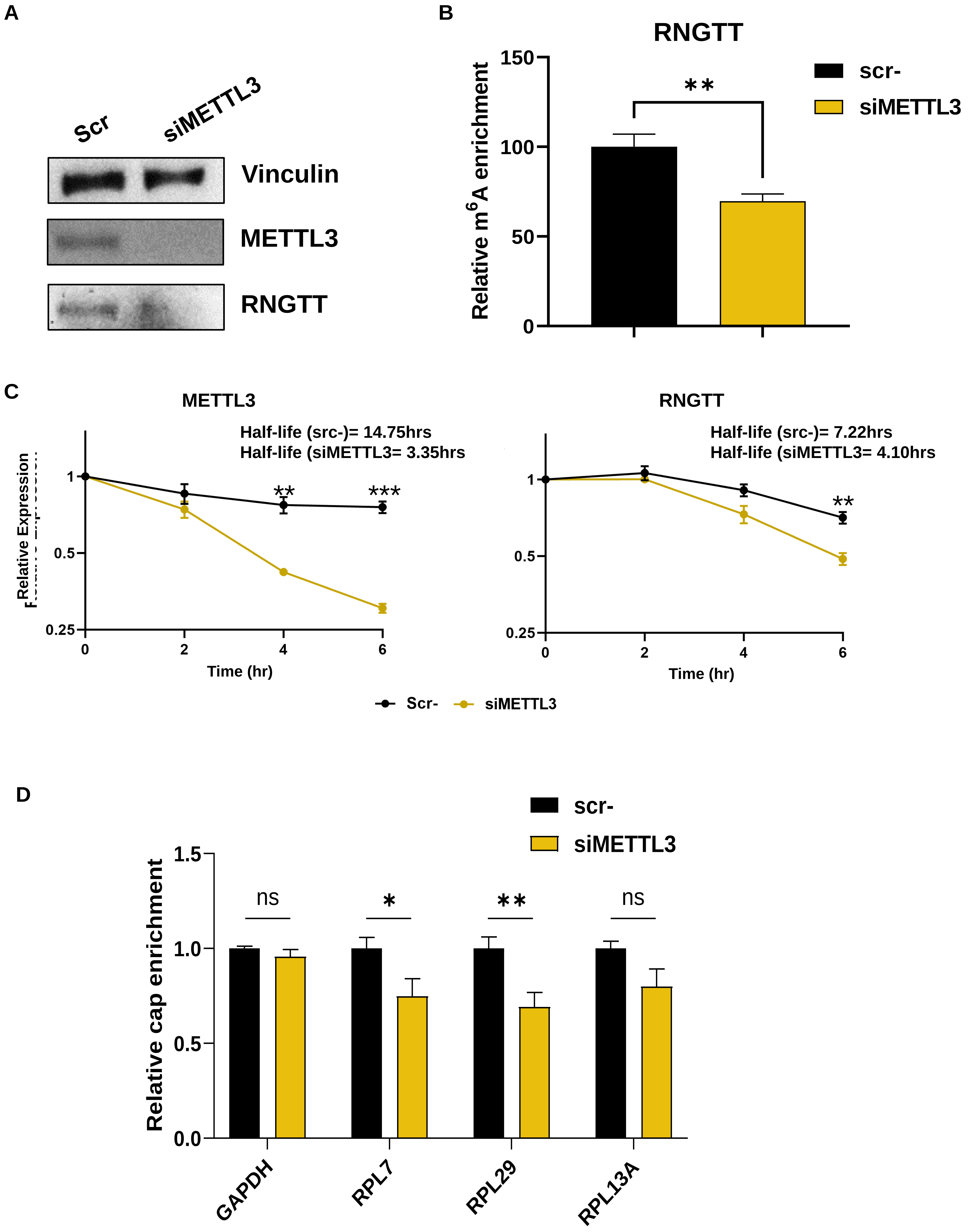
